# Macrophage-derived granulin drives resistance to immune checkpoint inhibition in metastatic pancreatic cancer

**DOI:** 10.1101/234906

**Authors:** Valeria Quaranta, Carolyn Rainer, Sebastian R. Nielsen, Meirion Raymant, Muhammad Shamsher Ahmed, Dannielle D. Engle, Arthur Taylor, Trish Murray, Fiona Campbell, Daniel Palmer, David A. Tuveson, Ainhoa Mielgo, Michael C. Schmid

## Abstract

The ability of disseminated cancer cells to evade the immune response is a critical step for efficient metastatic progression. Protection against an immune attack is often provided by the tumour microenvironment that suppresses and/or excludes cytotoxic CD8^+^ T cells. Pancreatic ductal adenocarcinoma (PDAC) is a highly aggressive metastatic disease with unmet needs, yet the immuno-protective role of the metastatic tumour microenvironment in pancreatic cancer is not completely understood. In this study we find that macrophage-derived granulin contributes to cytotoxic CD8^+^ T cell exclusion in metastatic livers. Mechanistically, we find that granulin expression by macrophages is induced in response to colony stimulating factor-1. Genetic depletion of granulin reduces the formation a fibrotic stroma, thereby allowing T cell entry at the metastatic site. While metastatic PDAC tumours are largely resistant to anti-PD-1 therapy, blockade of PD-1 in granulin depleted tumours restored the anti-tumour immune defence and dramatically decreased metastatic tumour burden. These findings suggest that targeting granulin may serve as a potential therapeutic strategy to restore CD8^+^ T cell infiltration in metastatic PDAC, thereby converting PDAC metastatic tumours, which are refractory to immune checkpoint inhibitors, into tumours that respond to immune checkpoint inhibition therapies.

## Introduction

The ability of the immune system to identify and destroy cancer cells is a primary defence mechanism against cancer (1). Accordingly, almost all solid cancers contain inflammatory immune cell infiltrates (2). CD8^+^ cytotoxic T cells, also known as cytotoxic T cells (CTLs), are key effectors of the immune response against cancer (1,3) and their presence in tumours is associated with a good clinical outcome in many tumour types, including ovarian, colon, breast, and pancreatic cancer (4-6). Paradoxically, although effector T cells have the capacity to identify and destroy cancer cells, other immune cells, particularly cells from the innate immune system such as macrophages, can have the opposite effect and promote tumour progression (7,8), resistance to therapy (9,10) and metastasis (11,12), thus, tumour-associated inflammation is considered a hallmark of cancer (13).

The importance of this dual function of the immune system in oncogenesis is demonstrated by the clinical success of anti-cancer immunotherapies, particularly the success achieved by the inhibition of immune checkpoint receptors (1,14). Indeed, the use of immune checkpoint inhibitors has recently been shown to be beneficial for many types of cancers, with anti-programmed cell death protein 1 (PD-1) inhibitors being one of the leading candidates (15). However, immune checkpoint inhibitors only work if effector T cells are infiltrated in tumours. Pancreatic tumours are particularly poorly infiltrated by effector T cells and thus, inhibition of immune checkpoint receptors alone did not show any benefit in pancreatic cancer patients (16,17). Pancreatic cancer is characterized by a rich and desmoplastic tumour stroma, also called tumour microenvironment (TME), identified by high numbers of activated fibroblasts, collagen deposition, and extensive myeloid cell infiltration, which all together critically impact the disease progression and its response to therapy (18). The TME is also thought to be a major barrier to CD8^+^ T cell infiltration in pancreatic tumours (19-21) and it is necessary to overcome this immune/fibrotic-protective barrier for the successful use of immune checkpoint inhibitors (21,22).

In the TME, macrophages represent a major component of the tumour infiltrating cells and they are implicated in various TME-mediated mechanisms that enable cancers to evade the immune response by restricting local T cell infiltration and T cell effector functions (21,23,24). Macrophages are innate immune cells with a high plasticity and depending on the activation signals, macrophages can acquire a spectrum of phenotypic states. In respect to cancer, macrophages can be polarized into M1 -like inflammatory macrophages that will activate a tumoricidal immune response (hereafter also referred to as M1-like), or into anti-inflammatory, immunosuppressive macrophages (hereafter also referred to as M2-like) that potently suppress other anti-tumour immune effector cells and thereby promote tumour progression (23). Indeed, a high density of macrophages, especially those exhibiting an immunosuppressive M2-like phenotype, correlates with poor clinical outcome in most human cancers (24,25). Accordingly, inhibition of myeloid cell recruitment into tumours have resulted in increased CD8^+^ effector T-cell infiltration and decreased tumour burden in mouse models and early-phase clinical trials (21,23,26-30). Yet, the mechanisms by which macrophages regulate T cell infiltration is only beginning to emerge.

Pancreatic ductal adenocarcinoma (PDAC) is a very aggressive metastatic disease and the fourth leading cause of cancer-related death. Currently, surgical resection is the best treatment option for PDAC patients, but, unfortunately, by the time PDAC is diagnosed, the majority of patients (∼ 80%) present with non-resectable metastatic cancer. Moreover, more than 60% of the patients whose tumours are removed, relapse with distant hepatic recurrence within the first 24 months after surgery (31-33). Thus, a better understanding of the mechanisms underlying the metastatic process in pancreatic cancer is critical to improve treatment and patient survival. We and others have recently identified that a desmoplastic TME also exists at the metastatic site in PDAC, which is mainly the liver, and that this fibro-inflammatory reaction is required for metastatic growth (11,34). However, whether and how the metastatic TME affects cytotoxic CD8^+^ T cell infiltration and function in the metastatic liver remains unexplored. Here, we found that macrophage-derived granulin is a key protein that supports CD8^+^ T cell exclusion and resistance to anti-PD-1 treatment. In fact, we show that depletion of granulin converts PDAC metastatic tumours, which are refractory to anti-PD1 treatment, into tumours that respond to anti-PD-1. Our findings provide pre-clinical evidence that support the rational for targeting granulin in combination with the immune checkpoint blocker PD-1 for the treatment of metastatic PDAC.

## Material and Methods

### Cells

Murine pancreatic cancer cells KPC FC1199, from here on referred as KPC, were generated in the Tuveson lab (Cold Spring Harbor Laboratory, New York, USA) isolated from PDA tumour tissues obtained from Kras^G12D/+^; p53^R17H/+^; Pdx1-Cre mice of a pure C57BL/6 background as described previously with minor modification^59^. Murine C57BL/6 Panc02 pancreatic ductal carcinoma cells were obtained from the NCI DCTD Tumor Repository, NIH. Panc02^luc/zsGreen^ or KPC^luc/zsGreen^ cells were generated by using pHIV Luc-zsGreen (gift from B. Welm, University of Utah, USA, Addgene plasmid no.39196) lentiviral particle infection. Infected cells were selected for high zsGreen expression levels using flow cytometry cell sorter (ARIA, BD). All cells were routinely tested negative for the presence of mycoplasma contamination. None of the cell lines used in this manuscript is listed in the ICLAC and NCBI Biosample database of misidentified cell lines.

### Mice

6-8 weeks old female C57BL/6 mice were purchased from Charles River. Grn^−/−^ mice (B6(Cg)- *Grn^tm1.1Aidi^*/J) and tdTomatoRed mice (B6.129(Cg)-Gt(ROSA)26Sor^tm4(ACTB-tdTomato-EGFP)Luo^/J) both on the C57BL/6 genetic background were purchased from The Jackson Laboratory. Kras^G12D/+^; p53^R172H/+^; Pdx1-Cre mice were purchased from CRUK, Cambridge Research Institute, Cambridge.

All animal experiments were performed in accordance with current UK legislation under an approved project licence PPL 40/3725 (Dr M Schmid). Mice were housed under specific pathogen-free conditions at the Biomedical Science Unit at the University of Liverpool. In general, for animal studies the group size was calculated by power analysis using a significance level kept at 5% and the power at 80% (according to approved corresponding Home Office Project License Application). For tumour studies, female animals age 6-8 weeks were used. Animals were randomly assigned to experimental groups. The investigators were not blinded to allocation during experiments and outcome assessments.

### *In vivo* animal studies

Liver experimental metastasis was performed by implanting 1×10^6^ KPC ^luc/zsGreen^ or Panc02 ^luc/zsGreen^ in 25ul PBS into the spleen of immunocompetent isogenic C57BL/6 mice using a Hamilton 29 G syringe as previously described ^8, 14^. For selected experiments, macrophages were depleted using CSF-1 neutralizing antibody (BioXCell, clone 5A1,) or CSF-1R inhibitor (Selleckchem, BLZ945). Anti-CSF-1 antibody was administered via intraperitoneal (i.p.) injection every 5 days with the first injection containing 1 mg and subsequent injections containing 0.5 mg. Rat IgG1 (BioXCell, clone HRPN) was used as isotype control. CSF-1R inhibitor BLZ945 was administered daily at a concentration of 200 mg/kg in 20% Captisol (Ligand Pharmaceuticals) by oral gavage. 20% Captisol was used as vehicle control. For immune checkpoint blockade, PD-1 antagonist (BioXCell, clone RMP-1) or Rat IgG2 (BioXCell, clone 2A3) isotype control were administered every 3 days by i.p. injection at 250 ug/dose. For T-cell depletion study, anti-CD8 (BioXCell, clone 2.43, 200ug/dose) was injected by i.p. 2 days prior intrasplenic implantation of KPC^luc/zsGreen^ cells into mice; at the day of KPC^luc/zsGreen^ implantation, and followed by injections every 4 days for the duration of the experiment. As isotype control, Rat IgG2 (BioXCell, clone LTF-2; 200ug/dose) was used. Treatment trials were initiated when transplanted tumours reached a mean volume of approximately 40 mm^3^ and 180 mm^3^ 7 and 14 days after implantation, respectively. Time points for treatment analysis were preselected based on the expected window of efficacy (after 1-2 weeks of treatment).

### Metastatic tumour burden quantification

Liver metastatic tumour burden was assessed, both *in vivo* and *ex-vivo*, by measuring bioluminescence signal (IVIS, Perkin Elmer) generated by KPC^luc/zsGreen^ or Panc02 ^luc/zsGreen^ cells. Bioluminescence signal was detected by intraperitoneal injection (i.p.) of Beetle luciferin (Promega; 3mg/mouse) and quantified as total flux (photons/sec). For some experiments, change in tumour volume in response to inhibitory Abs or drug treatment was assessed by 9.4 Tesla (T) horizontal bore Biospec MRI scanning (Bruker Biospin, Inc.). Mice livers were scanned *in vivo* before and after treatment using a T2-TurboRARE acquisition protocol. MRI images were acquired in coronal plane with 0.5 mm thickness and 0.1 mm spacing between each slice. A set of 23 slices was acquired to monitor the entire liver volume. Metastatic tumour volume was quantified by Image J software. Percentage change in metastatic tumour burden (before and after treatment) was obtained by calculating the sum of tumour area of all slices spanning the liver and multiplying it by 0.6 mm (interslice distance). The MRI acquisition used the following parameters: 2500 ms TR; 24 ms TE; 30×30 mm FOV; 240 × 240 image size. At indicated endpoints, liver metastatic lesions size and frequency were quantified based on haematoxylin and eosin staining of paraffin-embedded liver sections. Zeiss microscope and ZEN imaging software was used.

### Preparation of Conditioned media (CM)

Conditioned medium from Panc02, KPC cancer cells and bone marrow derived macrophages (BMM) was generated according to previous reports (35). Briefly, the medium was removed from 70% confluent cells and the cells were washed three times with PBS before addition of serum-free medium. Cells were incubated for 18-24h in serum-free medium and then collected and filtered through 0.45 μm filters before use.

### Assessment of *Granulin* gene expression in *in vitro* culture Bone Marrow derived Macrophage (BMM)

Primary murine macrophages were generated by flushing the bone marrow from the femur and tibia −1 of C57BL/6 mice followed by incubation for five days in DMEM containing 10% FBS and 10ng ml^-1^ murine M-CSF (Preprotech). BMM were subsequently stimulated with DMEM containing 2 % serum in the presence or absence of murine recombinant M-CSF (Preprotech) for 24 hour. Alternatively, BMM were stimulated with KPC or Panc02 CM for 24 hours in the presence of absence of anti-CSF-1 inhibitory antibody ([2.5 μg/ml; BioXCell). BMMs were finally lysed in RLT buffer + β-Mercaptoethanol and *Granulin* expression was assessed by qPCR.

### ELISA

Conditioned media from Panc02, KPC cancer cells and BMMs was obtained to measure the production of murine M-CSF by Quantikine ELISA kit (R&D System) according to the manufacturer’s instruction.

### T cell adoptive transfer experiment

For T cell adoptive transfer, experimental metastasis was induced by intrasplenic implantation of 1×10^6^ KPC^luc/zsGreen^ cells into tdTomatoRed+, WT, and Grn^−/−^ mice. After 13 days, tumour bearing tdTomatoRed+ mice were euthanized and spleens were dissected to isolate CD8^+^ T cells (Miltenyi, CD8a^+^ T cell isolation Kit). Isolated CD8^+^ T cells were stimulated with Dynabeads Mouse T- Activator CD3/CD28 (Life Technology) following manufacturer’s instruction, and incubated for 24 hours at 37°C. The next day, 1.5×10^6^ tdTomoatoRed^+^ CD8^+^ T cells were injected into the tail vein of tumour bearing WT and Grn^−/−^ mice. After 24 hours, mice were sacrificed and livers were collected and embedded in Optimal Cutting Temperature (OCT) medium. 5 μm liver sections were stained for DAPI (Life Technology, 1:500) and spatial localization of adoptive transferred dtTomatoRed^+^ CD8+ T cells was assessed by fluorescence microscopy measuring dtTomato Red^+^ signal.

### Bone marrow transplantation

Bone marrow transplantation was performed by reconstituting the bone marrow of lethally irradiated (10 Gy) female, 6-week-old C57BL6 mice by tail vein injection of 5 × 10^6^ total bone marrow cells isolated from Grn^−/−^ mice or WT mice ^14^. After 4 weeks, engraftment of Grn^−/−^ bone marrow was assessed by genomic DNA PCR according to The Jackson Laboratory protocol on peripheral blood cells from fully recovered bone-marrow-transplanted mice. After confirmation of successful bone marrow reconstitution, mice were enrolled in tumour studies.

### Flow cytometry

Single cell suspensions from murine livers were prepared by mechanical and enzymatic disruption in Hanks Balanced Salt Solution (HBSS) with 1mg/mL Collagenase P (Roche) as previously described 14. Briefly, cell suspensions were centrifuged for 5 minutes at 1200 rpm, resuspended in HBSS and filtered through a 500 μm polypropylene mesh (Spectrum Laboratories). Cell suspensions were resuspended in 1ml 0.05 % trypsin and incubated at 37° C for 5 minutes. Cells were filtered through a 70 μm cell strainer and resuspended in PBS + 0.5% BSA. Cells were blocked for 10 minutes on ice with FC block (BD Pharmingen, Clone 2.4 G2,) and then stained with Sytox-blue viability marker (Life Technologies) and conjugated with antibodies against CD45 (Clone 30F-11), F4/80 (Clone BM8), CD8 (Clone 53-6.7), CD206 (Clone C068C2), PD-1(Clone 29F.1A12), CD69 (Clone H1.2F3), IFNγ (Clone XMG1.2), Ki67 (Clone 16A8); Ly6G (Clone 1A8) all purchased from Biolegend; Granzyme B (eBioscience, clone NGZB). For intracellular staining, cells were first fixed (eBioscience, IC fixation buffer) and permeabilized (eBioscience, 1× permeabilization buffer).

To assess IFNγ expression levels in metastasis derived CD8^+^ T cells, magnetically isolated CD8a^+^ T cells from metastatic livers were stimulated with 50 ng/ml phorbol 12-myristate 13- acetate (PMA) (Sigma Aldrich) and 1 μg/ml of Ionomycin (Sigma Aldrich) for 5 hours at 37° C in the presence of Brefeldin A (eBioscience, 1:100) and subsequently stained for IFNγ.

Flow cytometry was performed on a FACS Canto II (BD Bioscience) and FACS cell sorting was carried out using FACS Aria (BD Bioscience). Cells were sorted directly in RLT buffer + β- Mercaptoethanol accordingly with the manufacturer’s instruction for RNA isolation (Qiagen).

### Magnetic beads isolation of cells

Samples for magnetic bead isolation were prepared from livers as described above for preparation of flow cytometry samples. Samples were stained and CD8a^+^ or F4/80^+^ cells were isolated according to the manufacturer’s instruction (Miltenyi).

### *In vitro* T-cell activation assay

Primary splenocytes were obtained from spleens of naïve C57BL/6 mice. Dissected spleens were dissociated in MAC buffer and passed through a 70μm cell strainer to obtain a single cell suspension. Cells were centrifuged (400 × g) and red blood cells were lysed using 1× Red blood lysis buffer (Biolegend). Obtained splenocytes were cultured in RPMI supplemented with 10% FBS. For T cell activation assays, splenocytes were stimulated using Dynabeads Mouse T- Activator CD3/CD28 (Life Technology). Activated splenocytes (S) were then co-cultured with macrophages (M) magnetically isolated cells from day 6 and day 14 metastasis bearing livers (4:1 ratio, S: M). Cells were plated in 96 well plates and incubated at 37°C for 24h. Subsequently, Brefeldin A (eBioscience, 1:100) was added to the cells for 5h. Cells were then harvested and stained with CD8 (Biolegend, clone 53-6.7) and IFNγ (Biolegend, Clone XMG1.2) antibodies and analysed by flow cytometry.

### *In vitro* T- cell proliferation Assay

For T cell proliferation assay, splenocytes derived from naïve C57BL/6 mice were labelled with 5 μ? Carboxyfluorescein Diacetate Succinimidyl Ester (CFSE) (Biolegend) and incubated for 10 minutes at 37°C in the dark. Cells were then resuspended in RPMI 1640 supplemented with 10% FBS and stimulated with Dynabeads Mouse T-Activator CD3/CD28. Activated splenocytes (S) were then cocultured with macrophages (M) magnetically isolated cells from day 6 and day 14 metastasis bearing livers (4:1 ratio, S: M). Cells were plated in 96 well plates and incubated at 37°C for 72 hours. Subsequently, cells were harvested and stained with CD8 antibody (Biolegend, clone 53-6.7). Proliferating CD8^+^ T cells were tracked by flow cytometry.

### RT-qPCR

Total RNA purification was performed with the RNeasy kit (Qiagen) and cDNA was generated using QuantiTect Reverse Transcription kit (Qiagen) according to the manufacturer’s instructions. 500 ng of total RNA was used to generate cDNA. Quantitative polymerase chain reaction (qPCR) was performed using 5x HOT FIREPol EvaGreen qPCR Mix Plus (ROX) (Solis Biodyne) on an MX3005P instrument (Stratagene). Three-step amplification was performed (95° C for 15 seconds, 60° C for 20 seconds, 72° C for 30 seconds) for 45 cycles. Relative expression levels were normalized to *Gapdh* expression according to the formula 2^Λ^–(Ct *gene of interest* – Ct *Gapdh).* Fold increase in expression levels were calculated by comparative Ct method 2^Λ^- (ddCt) (36).

The following QuantiTect Primers Assays were used to assess mRNA levels: *Gapdh* (Mm_Gapdh_3_SG; QT01658692), *Cxcl10* (Mm_Cxcl10_1_SG; QT00093436), *Cd86* (Mm_Cd86_1_SG; QT01055250), *Ifng* (Mm_Ifng_1_SG; QT01038821), *Il12* (Mm_Il12b_1_SG; QT00153643), *H2-Aa* (Mm_H2-Aa_1_SG; QT01061858), *Retnla* (Mm_Retnla_1_SG; QT00254359), *Tgfb* (Mm_Tgfb1_1_SG; QT00145250), *Il10* (Mm_Il10_1_SG; QT00106169), *Arginase* (Mm_Arg1_1_SG; QT00134288), *Gzmb* (Mm_Gzmb_1_SG; QT00114590), *Tnf* (Mm_Tnf_1_SG; QT00104006), *Prf1* (Mm_Prf1_1_SG; QT00282002), *Mrc1* (Mm_Mrc1_SG; QT00103012), *Granulin* (Mm_Grn_1_SG; QT01061634). All primers were purchased from Qiagen.

### Immunofluorescence

Murine liver tissues were fixed using a sucrose gradient method to preserve the zsGreen fluorescence. Briefly, livers were fixed in 4% Formaldehyde + 10% sucrose in PBS for 4 hours and then transferred to 20% sucrose in PBS for 8-10 hours. Tissues were transferred into 30% sucrose for an additional 8-10 hours, embedded in OCT medium and stored at −80°C.

For immunofluorescence staining 5 μm liver sections were permeabilized by 0.1% TritonX-100 (Sigma Aldrich) for 10 minutes. Unspecific bindings were prevented by using PBS +8% Normal goat serum for 1 hour at RT. Tissue sections were incubated overnight at 4 °C with the following antibodies: αSMA (Abcam, ab5694, 1:100); Relm-α (Abcam, 39626, 1:100); Cleaved- Caspase 3 (Cell Signalling, 9661, 1:100); CSF1R (Santa Cruz Biotechnology, clone c-20, 1:100). The next day, tissue sections were washed in PBS and stained with the secondary antibody goat anti-rabbit conjugated to AlexaFluo594 (Abcam, 1:500) and DAPI (Life Technologies, 1:500) for 1 hour at RT. Zs Green/Luciferase transfected cells were detected by their intrinsic signal. Sections were finally mounted using Dako Fluorescent Mounting Medium.

Immunofluorescence staining was also performed in some cases on tissue sections obtained from livers directly embedded in OCT. In this case tissue sections were fixed in ice-cold acetone for two minutes and permeabilized with 0.1% TritonX-100. Sections were washed and incubated overnight at 4°C with the primary antibodies: CD8 (Biolegend, clone 53-6.7, 1:100), F480 (Biolegend, clone BM8, 1:100). The next day, tissue sections were washed in PBS and stained with the secondary antibody goat anti-rabbit conjugated to AlexaFluo594 (Abcam, 1:500); goat anti-rat conjugated to AlexaFluo488 (Abcam, 1:500) and DAPI (Life Technologies, 1:500) for 1 hour at RT. All tissue sections were imaged using an Axio Observer Light Microscope with the Apotome.2 (Zeiss) and quantified using the Zen Software (Zeiss). Quantification of intrametastatic (IM) and peripheral (P) CD8^+^ T cells was performed using Nis Elements, Advanced Research software (Nikon). Peripheral area of metastatic lesions was defined as the outer 40 % of the total lesion area.

### Immunohistochemical analysis

Deparaffinization and antigen retrieval was performed using an automated DAKO PT-link. Paraffin-embedded human and mouse liver metastatic sections were immunostained using the DAKO envision+system-HRP. Tissue sections were incubated overnight at 4°C with primary antibodies: aSMA (Abcam, ab5694, 1:200); CD8 (Dako, Clone 144b, 1:100); Cytokeratin 19 (Abcam, ab53119, 1:100); Granzyme B (Dako, 1:50); PD-1 (CD279) (Abcam, ab52587, 1:100); CD3 (Abcam, ab16669, clone SP7, 1:100); Granulin (R&D Systems, MAB25571, 1:50); iNOS (Abcam, ab15323, 1:100); MHC II (Abcam, ab25333, 1:100); COX-2 (Cambridge bioscience, aa570-598, 1:100); Ym-1 (StemCell Technology, 60130, 1:200); CD206 (Abcam, ab8918, 1:100); Ly6G (Biolegend, Clone A18, 1:100). Secondary-HRP conjugated antibodies were incubated for 30 minutes at RT. Staining was developed using diaminobenzidine and counterstained with hematoxylin.

### Picrosirius red staining

Paraffin embedded murine liver samples were de-waxed and hydrated using a graded ethanol series. Tissue sections were then treated with 0.2% phosphomolybdic acid and subsequently stained with 0.1 % Sirius red F3B (Direct red 80)(Sigma Aldrich) in saturated picric acid solution for 90 minutes at RT. Tissues were then rinsed twice in acidified water (0.5% glacial acetic acid) (Sigma Aldrich) before and after the staining with 0.033 % fast green FCF (Sigma Aldrich). Finally, tissues were dehydrated in three changes of 100 % ethanol, cleared in xylene and mounted. Picrosirius red staining was quantified using Image J software.

### Human tissue samples

Paraffin embedded human tissue sections from control healthy subjects and advanced PDAC patients with liver metastasis were obtained from the Liverpool Tissue Bank, University of Liverpool, UK and approved by NRES Committee North West –Cheshire REC15/NW/0477. All samples were pathologically confirmed.

### Statistical analysis

Statistical analysis of experiments with two groups was performed using a two-tailed unpaired Student’s t-test with 95% confidence interval. Statistical analysis of experiments with more than two groups was performed using nonparametric analysis of variance (ANOVA) test with comparisons between groups using Bonferroni’s multiple comparison test. All statistical analysis were performed using GraphPad Prism software, p<0.05 was considered significant. Statistical significance is indicated in the figures as follows: ***, p<0.001; **, p< 0.01; *, p< 0.05; n.s, not significant. Quantification of results is expressed as mean ± SEM unless stated otherwise. Number of mice used for *in vivo* experiments is reported in the figure legend. For immunohistochemical and immunofluorescence analysis of human and murine tissues sections at least n ≥ 3 different visual fields from each tissue section were used.

## Results

### CD8^+^ T cell function and infiltration are lost during metastatic progression

The common route of metastasis of PDAC is to the liver (31,37). To investigate whether anti-tumour immunity might affect PDAC metastasis, we first analysed by immunohistochemistry (IHC) techniques liver biopsies from advanced metastatic treatment naïve PDAC patients, and healthy livers. In healthy livers CD8^+^ T cells were evenly distributed, at a low density (Fig. 1A). In human liver samples containing metastatic PDAC, we identified large metastatic lesions with high numbers of cytokeratin positive (CK^+^) tumour cells (CK^+^ rich) showing few infiltrating CD8^+^ T cells (Fig. 1A, B; Supplementary Fig. S1A). In contrast, smaller metastatic tumour deposits (CK^+^ poor) were rich in CD8^+^ T cells (Figs. 1A, B). Similarly, small spontaneous metastatic lesions generated in the genetically engineered mouse model of pancreatic cancer (Kras^G12D^; Trp53^R172H^; Pdx1-Cre mice, KPC) showed higher T cell infiltration compared to larger established lesions (Supplementary Fig. S1B).

**Figure 1:**
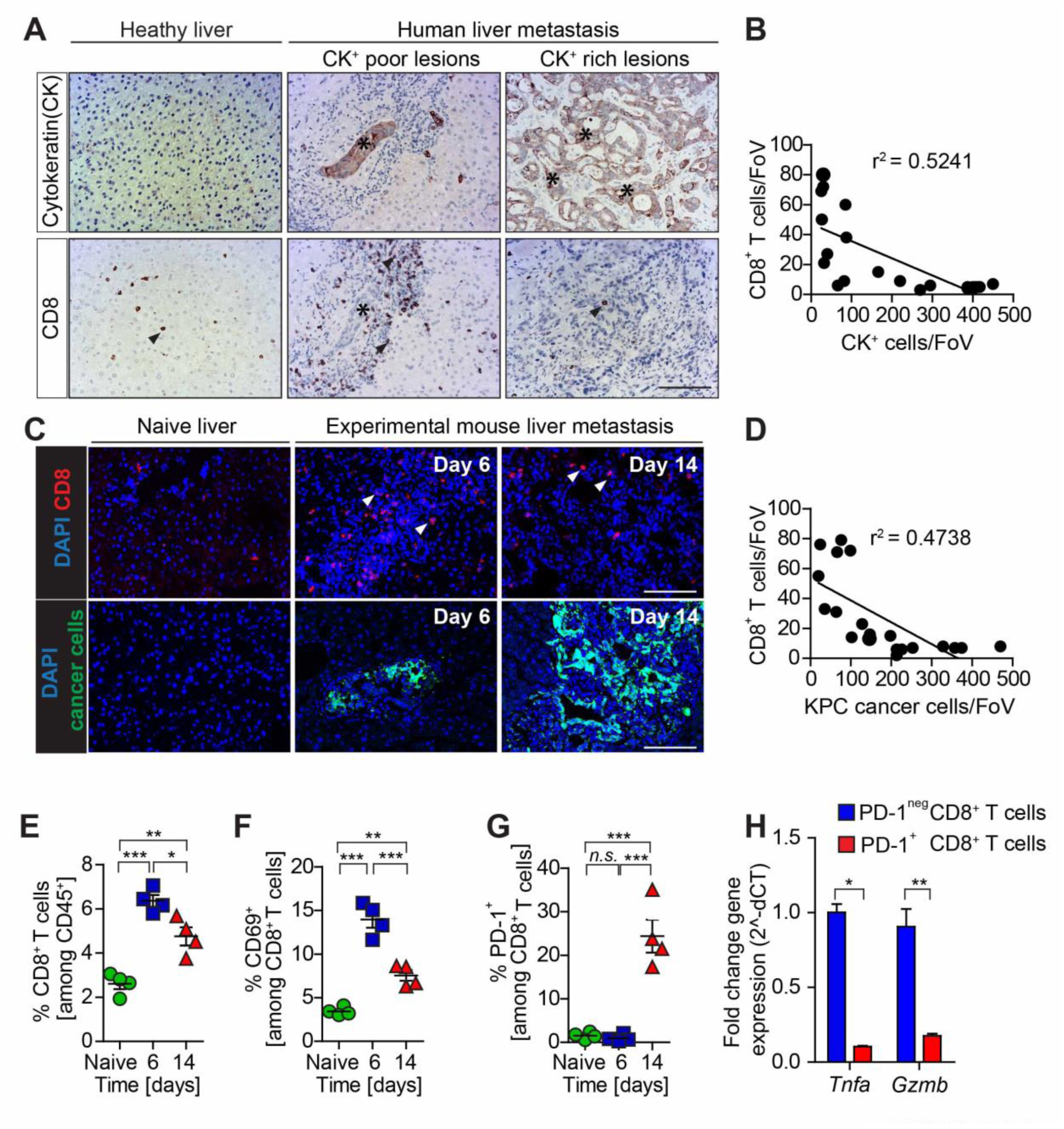
*Metastatic PDAC progression is accompanied by a loss of CD8^+^ T cell infiltration and activation*. (**A-B**) Identification of cytokeratin (CK)^+^ metastatic cancer cells and cytotoxic T cells (CD8^+^) infiltration by immunohistochemical analysis in human PDAC liver biopsies showing inverse correlation of CD8^+^ T cell and CK^^+^^ cancer cell numbers. (A) Representative micrographs for healthy liver, CK^^+^^ poor metastatic lesion, and CK^^+^^ rich metastatic lesion are shown. (B) Quantification of data. Metastatic cancer cells are indicated by asterisks. (n = 6 patients and n = 5 healthy subjects; n=3 different field of view (FoV)/patient were quantified in sequential tissue sections for CK^^+^^ and CD8^+^ cell staining, each dot represents FoV). (**C-D**) Immunofluorescence staining and quantification of metastatic KPC cells (zsGreen) and CD8^+^ T cell numbers in hepatic metastatic tumours resected 6 and 14 days after intrasplenic implantation of KPC^zsGreen/luc^ cancer cells. (C) Representative micrographs. (D) Quantification of data showing the inverse correlation of CD8^+^T cell and KPC^zsGreen/luc^ cell numbers. (n = 4 mice/time point, n = 5 FoV/mouse, each dot represents FoV) (**E-G**) Percentage of CD8^+^ T cells (E), CD69^+^ CD8^+^ T cells (F), and PD-1^+^ CD8^+^ T cells (G) cells over time (n= 4 mice / time point; individual data points, horizontal lines represent mean ± SEM) assessed by flow cytometry. (**H**) Quantification of *Tnfa* and *Gzmb* mRNA levels in metastasis infiltrating PD-1^+^ and PD-1^neg^ CD8^+^ T cells isolated by fluorescence activated cell sorting from established tumour bearing livers 14 days after intrasplenic implantation of 1×10^6^ KPC^luc/zsGreen^ cancer cells (n = 4 mice; mean ± SEM). Scale bars = 100μm; ***, *P*<0.001; **, *P* < 0.01; *, *P* < 0.05; n.s, not significant.

To further evaluate the changes in CD8^+^ T cell infiltration during PDAC metastasis, we next used a PDAC liver metastasis model in which KPC-derived pancreatic cancer cells expressing the dual reporter gene zsGreen and firefly luciferase (KPC^zsGreen/luc^) are intrasplenically injected and metastasize to the liver via the portal circulation (a common way of metastasis occurring in humans) (11,34,38). In agreement with the data obtained in PDAC patients and the autochthonous KPC model, we found that the metastatic infiltration of CD8^+^T cells was dependent on the number of cancer cells forming metastatic deposits in the liver. Small metastatic lesions mainly found at the early stage of metastatic progression (6 days post intrasplenic implantation) and characterized by low numbers of KPC cancer cells, showed high infiltration of CD8^+^ T cells in comparison to tumour free livers. In contrast, large metastatic lesions with abundant cancer cell numbers, mainly found at a later stage of metastasis progression (14 days post intrasplenic injection), showed a significant loss of CD8^+^ T cells (Figs. 1C, D). Multi-colour flow cytometry analysis of disaggregated murine livers further confirmed a significant increase in CD8^+^ T cells in day 6 small tumours, followed by a significant reduction in CD8^+^ T cell numbers in large tumours at day 14 (Fig. 1E).

The ability of CD8^+^ T cells to kill cancer cells depends on their activation state (39). To assess whether the identified CD8^+^ T cells were active, we next stained for the T cell activation marker CD69 and the inhibitory immune checkpoint receptor PD-1. Interestingly, CD8^+^ T cells from small tumours (day 6) were active (CD69^+^ and PD-1^+^), (Figs. 1F, G; Supplementary Fig. S1C), while CD8^+^ T cells isolated from large metastatic tumours (day 14) were inactive (CD69^-^ and PD-1+) (Figs. 1F, G). Indeed, gene expression analysis of the cytolytic factors *Tnfa* and *Gzmb* in metastasis infiltrating CD8^+^ T cells positive for PD-1 confirmed that the few infiltrating CD8^+^ T cells in large metastatic lesion (at day 14) are also exhausted (Fig. 1H). Collectively, these results indicate that CD8^+^ T cells are able to infiltrated small metastatic tumours but that during metastatic progression CD8^+^ T cell infiltration and cytotoxic function are lost.

### Immunosuppressive M2-like metastasis associated macrophages (MAMs) accumulate during PDAC metastasis

Tumour and metastasis associated macrophages can restrict CD8^+^ T cell effector functions depending on their activation state (22,40). Since large numbers of metastasis associated macrophages (MAMs) can be found at the hepatic metastatic site in PDAC (11), we next explored their activation state. Therefore, liver MAMs were identified as CD45^+^CD11 b^+^Ly6g^neg^Ly6C^dim/nes^ F4/80^+^ (Supplementary Fig. 2A) and gene expression of M1- and M2-like activation markers was assessed. MAMs isolated from small metastatic tumours (day 6) revealed a significant upregulation of immune stimulatory genes (*C-X-C motif chemokine 10, Cxcl10*; *interleukin 12, Il2; interferon gamma, Ifng*) and genes associated with antigen presentation (*H2Aa; Cd86),* resembling a pro-inflammatory M1-like phenotype (Fig. 2A). In contrast, macrophages isolated from large metastatic tumours (day 14) significantly upregulated the expression of immunosuppressive (transforming growth factor beta, *Tgfb, Arginase, Il10*) and anti-inflammatory M2-like markers (*mannose receptor C-type 1, Mrc1, resistin-like- α; Retnla*) (Fig. 2B). Histological analysis of small metastatic tumour tissue sections (day 6) confirmed a significantly higher number of myeloid-like cells expressing prototypical pro-inflammatory markers, including inducible nitric-oxide synthase (iNOS), major histocompatibility complex class II (MHC-II) and cyclooxygenase-2 (COX-2) (Fig. 2C). On the contrary, larger established metastatic tumours contained a higher number of myeloid-like cells expressing M2-like alternative activation markers, such as macrophage mannose receptor 1 (CD206) and Chitinase-like protein 3 (Ym-1) (Fig. 2D). Co- immunofluorescent staining further confirmed the accumulation of M2-like RELMα^+^ MAMs in close proximity to disseminated cancer cells (zsGreen) in large metastatic lesions (Fig. 2E).

**Figure 2:**
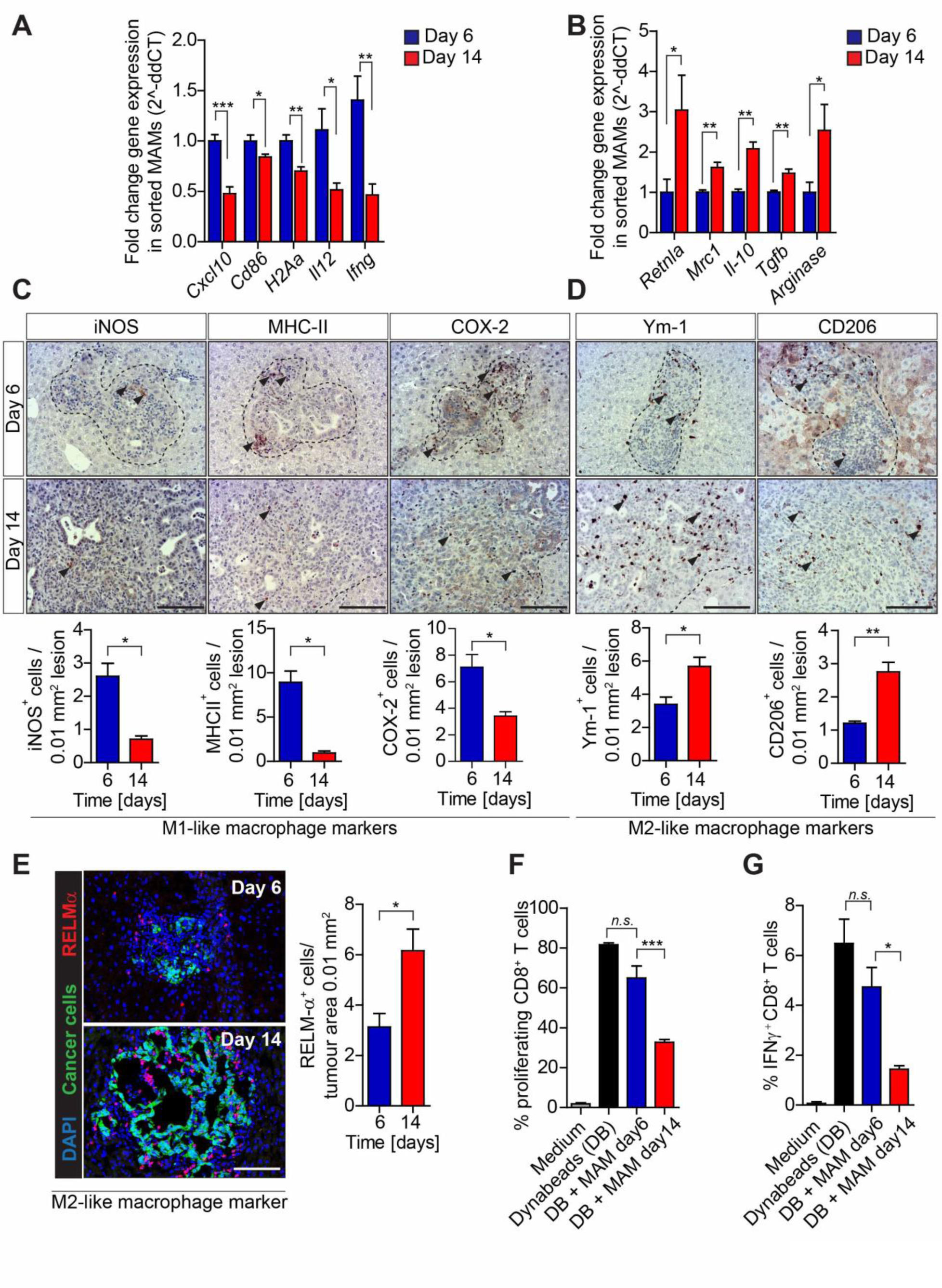
*Hepatic metastatic progression results in the accumulation of immune suppressive macrophages*. (**A-B**) qPCR analysis of multiple M1 and M2-macrophage associated genes in MAMs isolated from metastatic tumours resected 6 and 14 days after intrasplenic implantation of KPC ^luc/zsGreen^ cells (n = 4 mice / time point; mean ± SEM). (**C-D**) Representative images and quantification of immunohistochemical (IHC) analysis showing the number of cells staining positive for the M1 -like MAM markers (iNOS, MHC-II, COX-2) and M2-like MAM markers (Ym-1, CD206) in hepatic metastatic lesions from (A). (**E**) Representative immunofluorescent staining and quantification of M2-like macrophages (RELMa^+^) clustering around metastatic PDAC cells (zsGreen) in tumour bearing livers from (A). Nuclei were counterstained with DAPI. (**F-G**) Macrophages isolated from metastatic livers as described in A were assessed for their ability to (F) suppress splenic CD8^+^ T cell proliferation (CFSE dilution) (G) and activation (IFNγ expression levels) following CD3/CD28 Dynabeads (DB) stimulation (data are mean ± SD of 4 independent experiments). Scale bar= 100 μm; ***, P<0.001; **, *P* < 0.01; *, *P* < 0.05; n.s, not significant.

Since M2-like macrophages can execute potent inhibitory effects on cytotoxic CD8^+^ T cell functions (40), we next explored the impact of M1-like to M2-like MAM reprogramming on the apoptotic rate of disseminated tumour cells. We observed a higher apoptotic rate (cleaved caspase 3^+^) of cancer cells in small metastatic lesions (day 6) compared to large metastatic lesions (day 14) (Supplementary Fig. S2B). Finally, we assessed the ability of MAMs to suppress CD8^+^ T cells during metastatic progression. MAMs isolated from established metastatic lesions significantly suppressed CD8^+^ T cell proliferation (Fig. 2F) and activation *ex vivo* (Fig. 2G), compared to MAMs isolated from small metastatic lesions.

Together, these data suggest that metastatic progression in PDAC is accompanied by the reprogramming of MAMs towards an M2-like immunosuppressive phenotype that can inhibit cytotoxic CD8^+^ T cell functions.

### Pharmacological blockade of the CSF-1/CSF-1R axis reprograms MAMs towards an immune-stimulatory phenotype and restores CD8^+^ T cell mediated anti-tumour immunity in metastatic PDAC

Based on our findings, we reasoned that blocking the rewiring of M1 -like MAMs into immunosuppressive M2-like MAMs during metastatic progression could sustain tumoricidal CD8^+^ T cell functions and subsequently inhibit PDAC metastasis. Since colony stimulating factor 1 (CSF-1) can expand macrophage populations and drive M2 differentiation, CSF-1 and its cognate receptor CSF-1R are among the most advanced targets to inhibit tumour promoting macrophage functions in cancer (26,41). While CSF1/CSF-1R signalling has been extensively studied in primary tumours, its role in PDAC metastasis has not yet been explored. We found that CSF-1 is abundantly expressed by metastatic pancreatic cancer cells (Supplementary Fig. S2C) and that CSF-1R is expressed by MAMs in experimental and spontaneous PDAC metastasis models (Supplementary Figs. S2D, S2E). To assess whether the inhibition of CSF-1 affects MAM functions, T cell infiltration/activation, and metastatic PDAC progression, we induced hepatic metastasis by intrasplencially implanting KPC cells. Starting at day 7, after initial seeding has occurred (11) and small metastatic lesions are rich in CD8^+^ T cells (Fig. 1C), we administered neutralizing αCSF-1 monoclonal antibody for 2 weeks (Fig. 3A). αCSF-1 treatment reduced metastatic tumour burden (Fig. 3B). In respect to MAMs, αCSF-1 reduced not only overall MAM numbers (Fig. 3C), but specifically reduced the number of M2-like F4/80^+^CD206^+^ MAMs (Fig. 3D). Gene expression analysis of MAMs isolated by flow cytometry further confirmed that inhibition of CSF-1 induces a reprogramming of MAMs towards an inflammatory M1- like phenotype. Indeed, MAMs isolated from αCSF-1 treated mice showed a significant increase in the expression of pro-inflammatory cytokines and markers (*Cxcl10, Ifng, Il12, Cd86, H2-Aa*) whereas the levels of M2-like markers (*Mrc1, Retnla*) and immune suppressive factors (*Tgfb, Il10, Arginase*) were significantly reduced (Fig. 3E). Consistent with a rewired pro-inflammatory metastatic microenvironment, CD8^+^ T cell numbers and function were increased in αCSF-1 treated mice as shown by the increase in IFNγ and GzmB expression levels and an enhanced CD8^+^ T cell proliferation rate (identified by Ki67^+^CD8^+^ T cells) (Figs. 3F-I). It is noteworthy that αCSF-1 treatment also enhanced the proliferation rate (Ki67^+^) within PD-1^+^CD8^+^ T cells, suggesting a partial reinvigoration of exhausted PD-1^+^ CD8^+^ T cells within the metastatic niche (Fig. 3J). Importantly, tumour bearing mice treated with a small molecule inhibitor of CSF-1R (BLZ945) (42) showed a similar reduction in the metastatic tumour burden (Supplementary Figs. S3A-E) accompanied by an increase of pro-inflammatory M1 -like MAMs (Supplementary Figs. S3F, G) and CD8^+^ T cell in metastatic livers (Supplementary Fig. S3H) and an increase in apoptosis of metastatic cancer cells (Supplementary Fig. S3I) *in vivo.*

**Figure 3:**
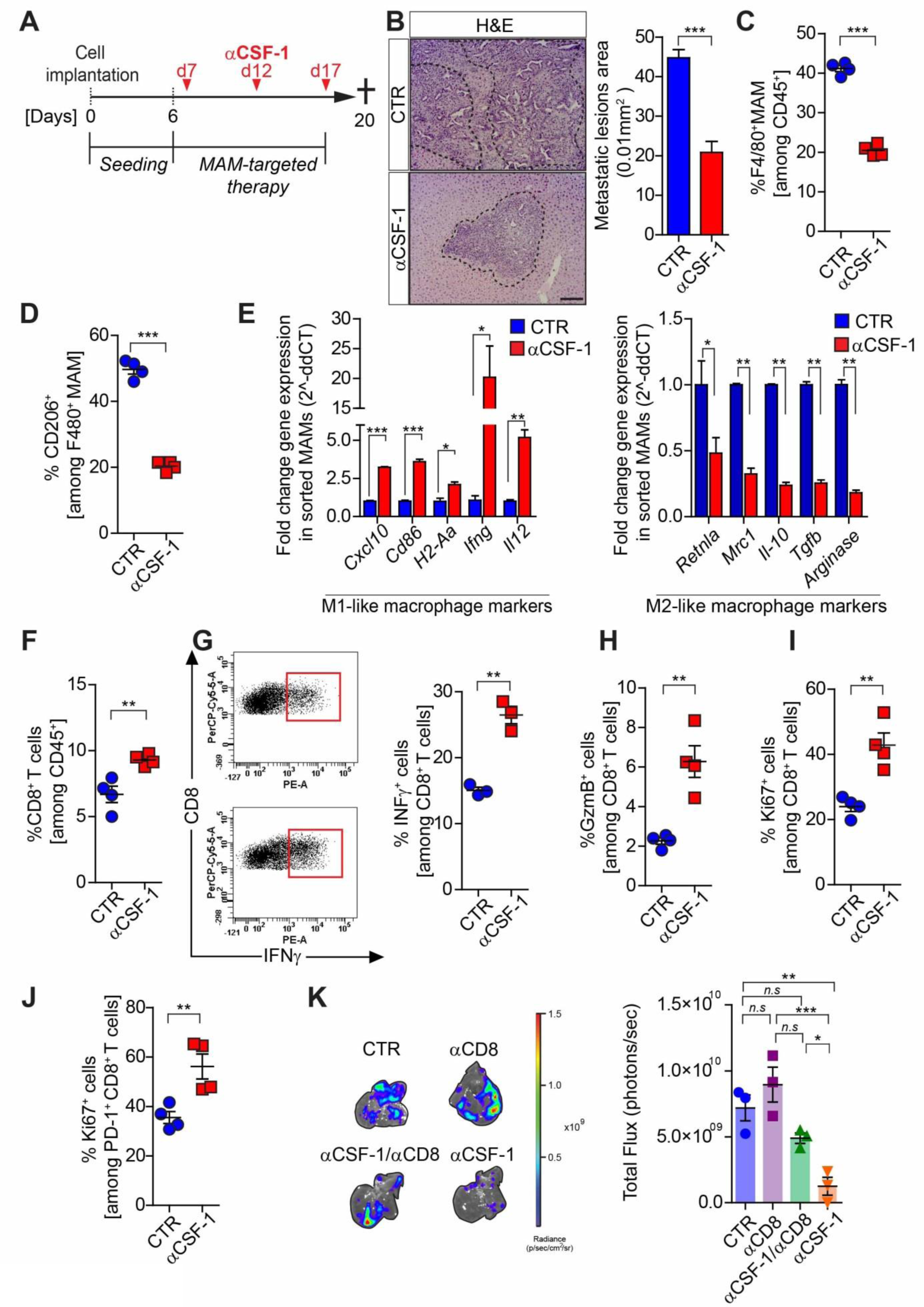
*Pharmacological blockade of CSF-1 reprograms MAMs and reinvigorates CD8^+^ T cell functions*. (**A-J**) Liver metastasis was induced by intrasplenic implantation of KPC^zsGreen/luc^ cells. After 7 days metastasis bearing mice were treated with IgG control (Ctr) or neutralizing αCSF-1 antibody. Mice were sacrificed at day 20. (**A**) Schematic illustration of the experiment. (**B**) Representative images and quantification of average metastatic lesion area in both treatment cohorts by H&E stained liver sections (n = 6 mice / group; mean ± SEM; two-tailed unpaired t-test). (**C-D**) FACS analysis quantifications of (C) total and (D) M2-like (CD206^+^) MAMs in tumour bearing livers (n= 4 mice / group; individual data points, horizontal lines represent mean ± SEM). (**E**) qPCR analysis of multiple M1 and M2-macrophage associated genes in MAMs isolated from tumour bearing livers at endpoint (n = 4 mice /group). (**F**) Metastasis infiltrating CD8^+^ T cells from tumour bearing livers in each cohort were evaluated by flow cytometry (n= 4 mice / group; individual data points, horizontal lines represent mean ± SEM). (**G**) Representative dot plot and quantification of data of IFNγ expression levels in metastasis infiltrating CD8^+^ T cells by flow cytometry (n = 3 mice /group, individual data points, horizontal lines represent mean ± SEM). (**H-J**) Quantification of (H) Granzyme B (GzmB), (I) proliferating (Ki67^+^) cells among total CD8^+^ T cells and (J) PD-1^+^ CD8^+^ T cells by flow cytometry (n = 4 mice /group, individual data points, horizontal lines represent mean ± SEM). (**K**) Liver metastasis was induced by intrasplenic implantation of KPC^zsGreen/luc^ cells in CD8^+^ T cells depleted (aCD8 Ab) or control (isotype IgG) mice. Cohorts were treated with αCSF-1 mAb or control IgG (Ctr) starting day 2. Representative images and quantification of *ex-vivo* metastatic tumour burden (total flux) by bioluminescence imaging at day 14 (n = 3 mice /group, individual data points, horizontal lines represent mean ± SEM). Scale bar = 100 μm; ***, *P*<0.001; **, *P* < 0.01; *, *P* < 0.05; n.s, not significant.

To further explore the role of macrophages in regulating cytotoxic CD8^+^ T cell infiltration and pancreatic cancer metastasis we next depleted CD8^+^ T effector cells alone or in combination with targeting macrophages (using αCSF1) and assessed metastatic tumour progression. Pharmacological depletion of CD8^+^ T cells abolished the anti-metastatic effect of αCSF-1 (Fig. 3K; Supplementary Figs. S4A, B) demonstrating that the ability of MAMs to support PDAC metastasis is in part due to their ability to suppress cytotoxic CD8^+^ T cells.

Taken together, our findings show that MAM targeted therapies can restore a pro-inflammatory environment in metastatic PDAC in which cytotoxic CD8^+^ T cells can infiltrate and kill metastatic cancer cells.

### CSF-1 inhibition reduces desmoplasia and sensitizes metastatic PDAC to anti-PD-1 treatment.

Since αCSF1 restores T cell infiltration in metastatic tumours (Fig. 3) we next evaluated whether combining αCSF1 with αPD1 treatment could reduce PDAC metastasis. To test this, we used a PDAC metastasis model (11) and treated tumours early, after initial seeding of cancer cells to the liver (Day 7), or later, at day 14, after T cell infiltration is lost (Fig. 1; Fig. 4A). We observed that in response to early intervention (day 7), metastatic tumour progression was significantly inhibited by both agents delivered as monotherapies (αPD-1= 49 +/− 3%; αCSF-1= 50 +/− 3%), but the anti-tumorigenic effect of αCSF-1 treatment was further potentiated when administrated in combination with αPD-1 (αCSF-1/αPD-1= 80 +/− 3%)(Fig. 4B). In contrast, in response to a later intervention on larger metastatic lesions (day 14), we observed that neither αPD-1 nor αCSF-1 alone was able to reduce metastatic tumour burden (KPC: αPD-1= 3 +/− 3%; αCSF-1= 22 +/− 2%; Panc02: αPD-1= 5+/−3%; αCSF-1= 21 +/− 4%) (Fig. 4B). In large metastatic lesions, only combinatorial treatment of αCSF- 1/αPD-1 was able to reduce metastatic tumour burden by 61+/− 4% (KPC) and 55+/− 3% (Panc02), respectively, in two independent PDAC cell lines (Fig. 4B).

**Figure 4:**
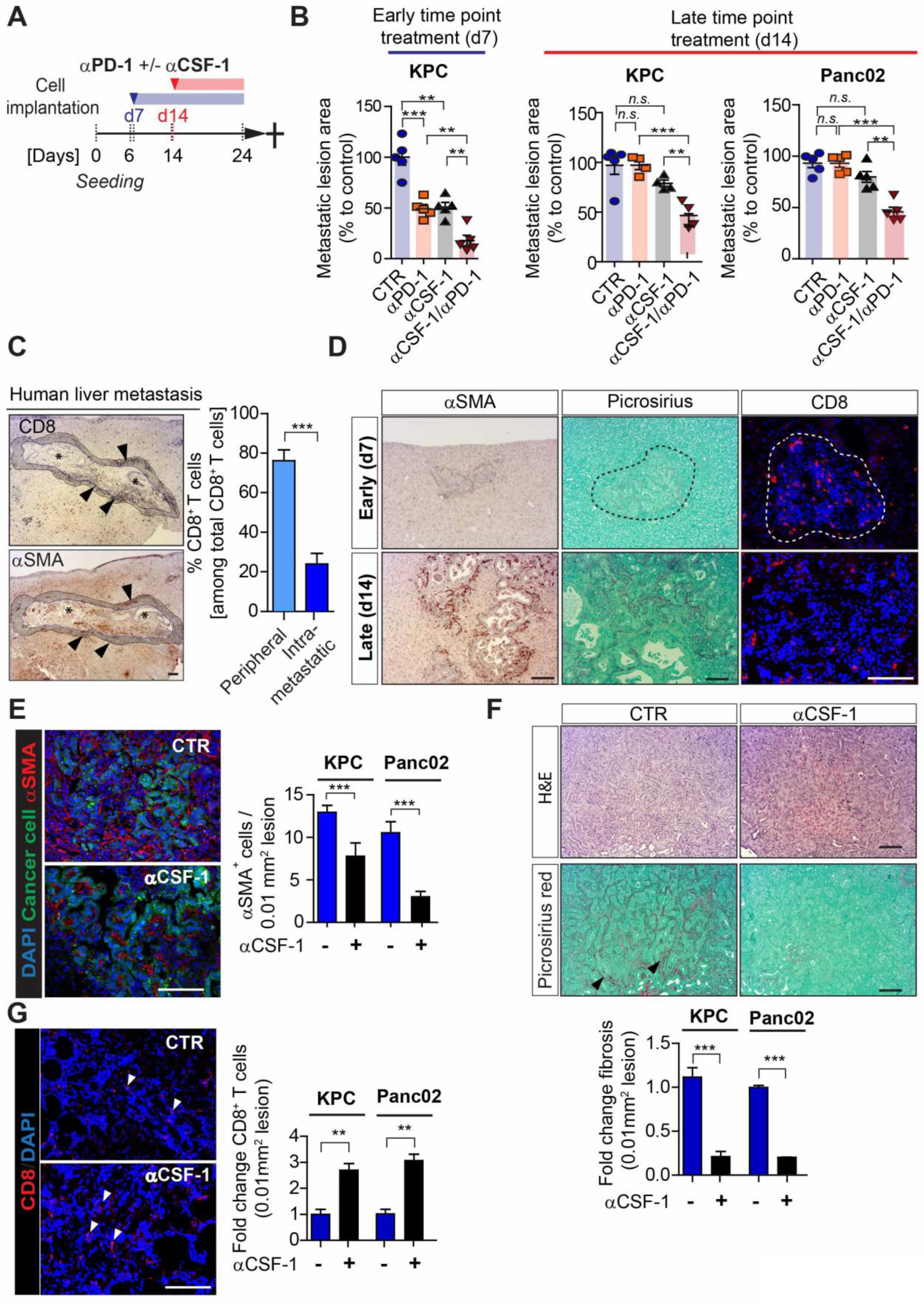
*CSF-1 inhibition reduces desmoplasia and sensitizes metastatic PDAC to anti-PD-1 treatment*. **(A-B)** Liver metastasis were induced by intrasplenic implantation of KPC^zsGreen/luc^ cells or Panc02^zsGreen/luc^. (A) Schematic representation of the treatment regimen conducted in metastatic mice. Cohorts were treated with control IgG Abs, αCSF-1 mAb alone, αPD-1 alone, or αCSF-1 and αPD-1 combined. Early time point treatment started at day 7, prior to T cell exclusion, while late time point treatment started at day 14. Tumour burden was quantified and analysed at day 24 (n = 4-5 mice / group). (B) Percentage of average change in metastatic lesion area compared to control in response to treatment assessed by haematoxylin and eosin (H&E) staining at endpoint (individual data points, horizontal lines represent mean ± SEM). (**C**) Representative immunohistochemical staining of T cells (CD8^+^) and myofibroblasts (αSMA^+^) in human PDAC metastatic livers and quantification of perimetastatic (P) and intrametastatic (IM) CD8^+^ T cells (n = 6 patients; mean ± SEM). (**D**) Liver metastasis was induced by intrasplenic implantation of 1×10^6^ KPC^zsGreen/luc^ cells. Livers were surgically removed 6 and 14 days later and assessed by αSMA^+^ (myofibroblasts), Picrosirius red (collagen deposition) and CD8^+^ T cell staining. (**E-G**) Representative micrographs and relative quantification of (E) myofibroblasts (αSMA^+^, red) cell frequency, (F) picrosirius red and H&E staining of sequential tumour sections showing area occupied by fibrotic stroma and (G) infiltrating CD8^+^ T cells, in liver tissue sections of mice treated at later time point of metastatic progression with IgG control or αCSF-1 ihnibitory antibody. Quantification of staining is refered to metastatic tumour generated by KPC ^zsGreen/luc^ or Panc02 ^zsGreen/luc^ pancreatic cancer cells implantation. Images are representative of KPC ^zsGreen/luc^ cancer cells derived liver metastatic lesions (n = 4-5 mice / group, mean ± SEM; additional treatment groups are shown in Supplementary Fig. S5).

Our results show that targeting MAMs with αCSF1 increases T cell infiltration in metastatic tumours and increases their response to αPD-1 treatment. In our previous work, we found that liver fibrosis develops in PDAC metastasis and that M2-like MAMs promote fibroblast activation in metastatic PDAC growth (11). Previous studies suggest that fibroblasts and collagen deposition can impair T cell infiltration in primary tumours (20,43). Thus, we hypothesize that αCSF1 treatment, which we know re-polarises M2-like macrophages into M1 -like macrophages (Fig. 3) may decrease fibroblast activation and collagen deposition, thereby supporting T cell infiltration and subsequently increase αPD-1 efficacy. To address this question we first assessed the spatial localisation of CD8^+^ T cells in advanced PDAC metastatic tumours from patients and mice. We found that advanced metastatic human tumours have low numbers of intra-metastatic CD8^+^ T cells and that the majority of CD8^+^ T cells present in these tumours accumulate at the periphery of metastatic lesions, where typically high numbers of myofibroblasts (αSMA^+^) are found (Fig. 4C). Similarly, metastatic murine tumours harvested at day 14 (large metastatic lesions), but not at day 7 (small metastatic lesions), showed a marked increase in αSMA^+^ myofibroblasts, collagen deposition (assessed by picrosirius red staining), that correlates with a decrease in CD8^+^ T infiltration (Fig. 4D).

Interestingly, in line with our hypothesis, we found that inhibition of CSF-1 altered the desmoplastic reaction at the metastatic site. In fact, KPC and Panc02 tumours treated with αCSF-1 alone or in combination with αPD-1 markedly reduced αSMA^+^ myofibroblast numbers (Fig. 4E; Supplementary Figs. S5D, H), collagen deposition (Fig. 4F; Supplementary Figs. S5A, C, F) and increased CD8^+^ T cells infiltration (Fig. 4G; Supplementary Figs. S5B, E, G) independently on the time of treatment intervention.

Taken together, these data suggest that the presence of a desmoplastic stroma affects cytotoxic CD8^+^ T cell infiltration in metastatic PDAC tumours, and that targeting macrophages using αCSF-1 inhibitor reduces the fibrotic reaction, increases T cell infiltration and response to αPD-1 therapy.

### CSF-1 inhibition reduces granulin expression in macrophages in vitro and in vivo.

We previously identified macrophage-derived granulin as a key effector protein for αSMA^+^ myofibroblast accumulation during PDAC metastasis (11). Thus, we next asked whether inhibition of CSF-1 could impair the expression of granulin in macrophages. Accordingly, we first assessed whether granulin expression in macrophages depends on CSF-1. Indeed, we found that recombinant CSF-1 is a strong inducer of granulin expression in primary bone marrow derived macrophages (BMM) (Fig. 5A). Moreover, pancreatic cancer cells, which abundantly secrete CSF-1 (Fig. 5B), induce the expression of granulin in macrophages in a CSF-1 dependent manner since adding a CSF-1 neutralizing antibody significantly downregulates granulin expression in primary BMM cultured in the presence of KPC or Panc02 conditioned media (CM) *in vitro* (Fig. 5C). In addition, we found a significant reduction of granulin expression in macrophages isolated from αCSF-1 treated metastatic tumours (Fig. 5D), which was further confirmed on protein level by IHC analysis of liver KPC and Panc02 metastatic tumour sections (Fig. 5E). Taken together, these results demonstrate that CSF-1 is necessary and sufficient to induce granulin expression in macrophages.

**Figure 5:**
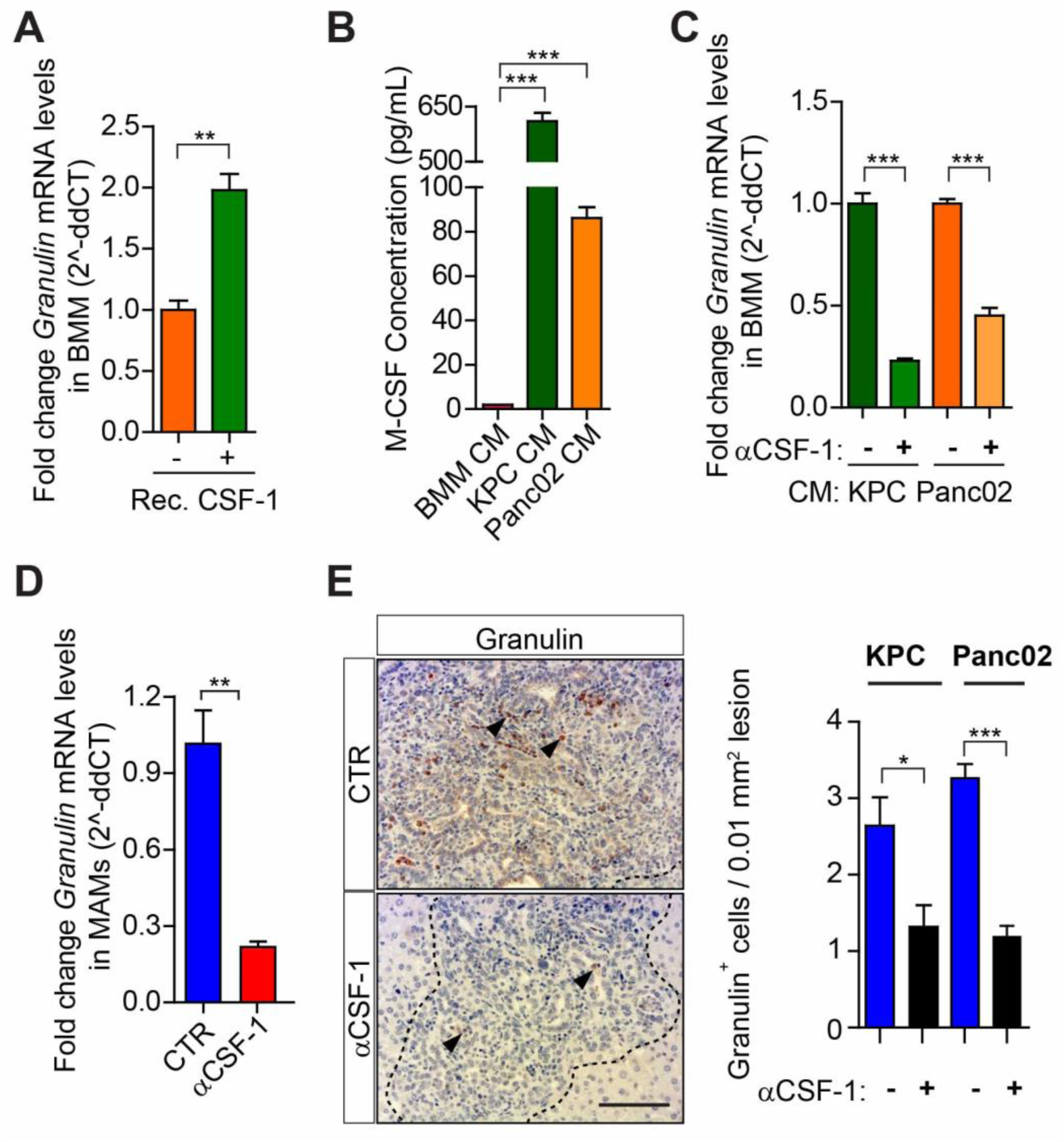
*CSF-1 is necessary and sufficient to induce granulin expression in macrophages*. (**A**) Quantification of *Granulin* mRNA levels by qPCR in primary unstimulated macrophages and macrophages exposed to recombinant CSF-1. (**B**) Quantification of CSF-1 protein levels in conditioned medium (CM) generated from bone marrow macrophages (BMM), and KPC and Panc02 pancreatic cancer cells using ELISA. (**C**) Quantification of *Granulin* mRNA levels by qPCR in primary unstimulated macrophages and macrophages exposed to KPC and Panc02 conditioned media (CM) in the presence of IgG control (Ctr) or neutralizing αCSF-1 mAb. (**D-E**) Mice were intrasplenically implanted with KPC^zsGreen/luc^ or Panc02 ^zsGreen/luc^ cells and treated with IgG control (Ctr) or neutralizing αCSF-1 mAb after 14 days. Livers were harvested from euthanized mice after 24 days. (D) Quantification of *Granulin* expression in MAMs isolated from metastatic lesions at endpoint (n = 4 mice / group; mean ± SEM). (E) Representative images and quantification of Granulin in metastatic livers by immunohistochemical analysis in KPC ^zsGreen/luc^ and Panc02 ^zsGreen/luc^ derived liver metastatic lesions (n = 6 mice / group; mean ± SEM).

### Genetic depletion of granulin restores CD8^+^ T cell infiltration in metastatic tumours, but T cell dysfunctionality remains.

Since granulin expression is triggered by CSF-1 and CSF-1 inhibition increases T cell infiltration in metastatic lesions, we next evaluated the effect of granulin depletion in T cell infiltration. To address this question, we intrasplenically injected KPC-derived cells into chimeric mice deficient of granulin (Grn^−/−^) in the bone marrow (BM) compartment (WT^+^ Grn^−/−^ BM mice). Intra-metastatic CD8^+^ T cell accumulation was markedly improved in granulin deficient mice (Figs. 6A, 6B) which as we previously described (11) are defective in myofibroblast activation (Figs. 6A, 6C). Since we previously showed that granulin depletion decreases metastatic tumour burden (11) and we know that smaller metastatic lesions have higher infiltration of cytotoxic T cells (Fig. 1), one could think that the effect of granulin depletion on T cell infiltration is a consequence of the reduction in the size of the metastatic lesions. To understand whether granulin depletion has a direct impact on CD8^+^ T cell infiltration we next performed an adoptive transfer of CD8^+^ T cells (derived from tdTomatoRed WT mice; tdTR) into WT mice or granulin deficient mice (Grn^−/−^) (Fig. 6D). Although overall metastatic tumour growth was significantly lower in granulin deficient mice (data not shown), livers of WT mice contained beside mainly large metastatic deposits also a few small tumour lesions. Thus, we compared CD8^+^ T cell numbers in metastatic lesions of equal sizes from both WT and Grn^−/−^ mice and found that depletion of granulin caused a significant increase of intra-metastatic (IM) tdTR CD8^+^ T cells in metastatic lesions of equal size compared to WT tumours while decreasing the number of peripheral (P) CD8^+^ T cells trapped at the tumour border (Fig. 6E). Noteworthy, depletion of granulin did not alter the overall number of CD8^+^ T cells in metastatic tumours (Fig. 6F), nor it improved their activation state as assessed by IFNγ expression levels (Fig. 6G).

**Figure 6:**
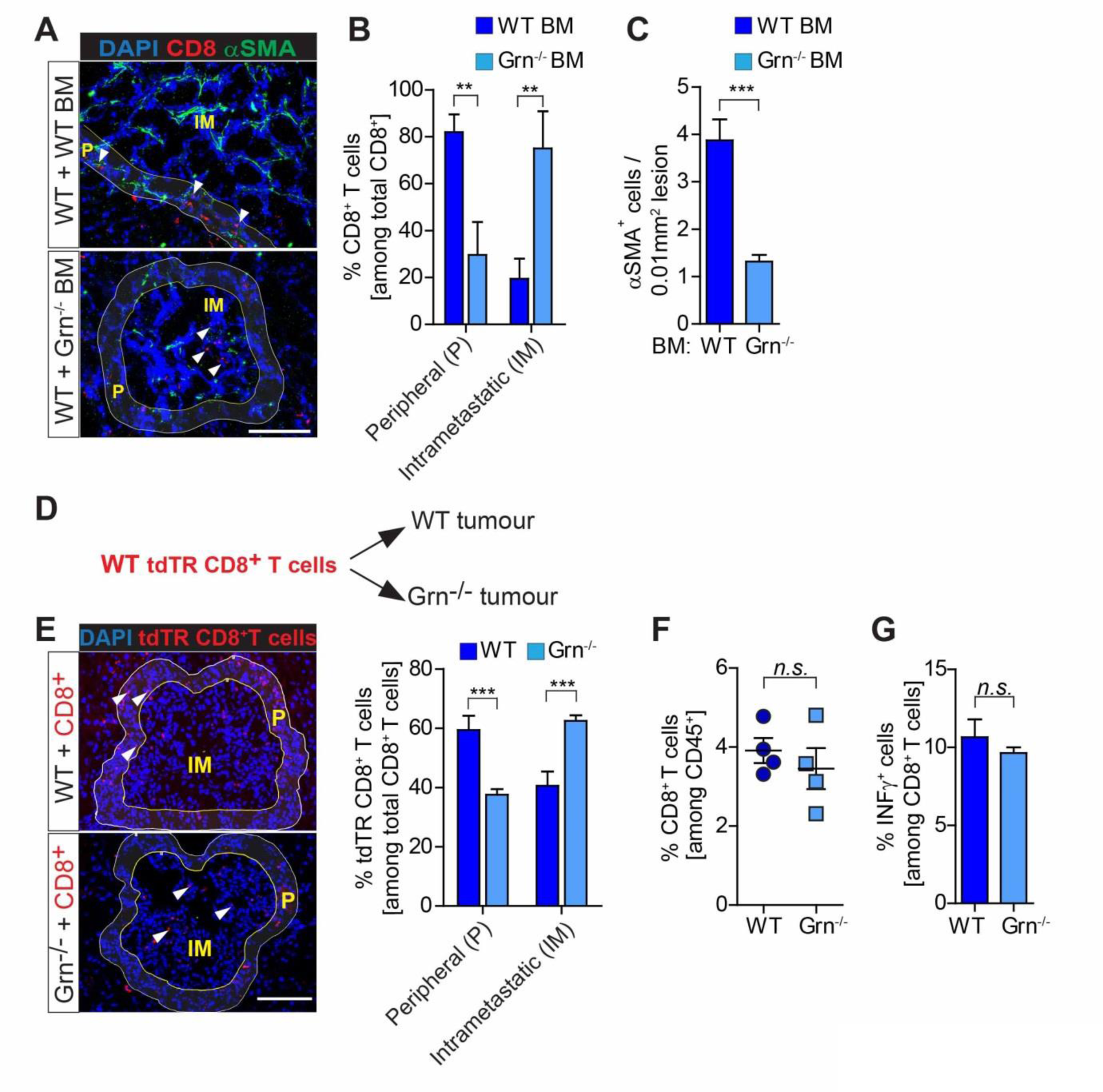
*Genetic depletion of granulin results in increased metastatic CD8^+^ T cell entry, but CD8^+^ T cells remain dysfunctional*. (**A-C**) Liver metastasis was induced by intrasplenically implantation of 5x10 KPC cells into chimeric WT^+^ WT BM and WT^+^ Grn^−/−^ BM mice. Entire livers were resected 14 days later and analysed. (A) Representative immunofluorescence staining of CD8^+^ T cells and αSMA^+^ myofibroblasts in metastatic livers. Intrametastatic (IM) and peripheral areas (P) are indicated. (B) Quantification of peripheral (P) and intrametastatic (IM) CD8^+^ T cells in metastatic livers. (C) Quantification of αSMA^+^ cells in metastatic livers (WT BM, n = 5; Grn^−/−^ BM, n = 6; mean ± SEM). (**D-E**) Liver metastasis was induced by intrasplenically implantation of KPC^zsGreen/luc^ cells into dtTR, WT and Grn^−/−^ mice. At day 13, isolated CD8^+^ T cells were isolated from tdTR mice and adoptively transferred into tumour bearing WT and Grn^−/−^ mice.24 hours later livers were resected and analysed for tdTR CD8^+^ T cell infiltration. (**D**) Schematic illustration of the experiment. (E) Representative immunofluorescence images and quantification of data showing percentage of tdTR CD8^+^ T cells (red) found in perimetastatic (P) and intrametastatic (IM) regions upon adoptive transfer (n = 3 mice / group; mean ± SEM). (**F-G**) FACS analysis quantifications of (F) CD8^+^ T cell number and (G) CD8^+^ T cell activation (IFNγ^+^ CD8^+^ T cell) in hepatic metastatic tumours resected 14 days after intrasplenic injection of KPC cells into WT and Grn^−/−^ mice. Scale bar = 100 μm. *P*<0.001; **, *P* < 0.01; *, *P* < 0.05; n.s, not significant.

Taken together, our results suggest that depletion of granulin improves CD8^+^ T cell infiltration into metastatic tumours, but does not restore their function.

### Depletion of granulin restores the response of metastatic PDAC to PD-1 therapy.

We queried whether the observed increase in CD8^+^ T cell entry into granulin deficient metastatic tumours, might then also improve their response to αPD-1 therapy. To address this question, we induced liver metastasis in immuno-competent WT and Granulin deficient (Grn^−/−^) mice by intrasplenic implantation of KPC^zsGreen/luc^ cells. Starting at day 14, when metastatic lesions are large, poorly infiltrated by CD8^+^ T cells and rich in myofibroblasts and collagen (Fig. 4D) (11), mice were treated with either αPD-1 or IgG control antibody. To assess the response of metastatic tumours to treatment, metastatic tumour burden was quantified by *in vivo* bioluminescence imaging (BLI) techniques at day 14 prior treatment and at endpoint (day 24) (Fig. 7A). In agreement with our previous observations, single agent αPD-1 therapy did not show any efficacy in metastatic tumour bearing WT mice (Fig. 7B, C). In contrast, αPD-1 treatment in granulin depleted mice (Grn^−/−^) caused a dramatic decrease in metastatic progression, with even partial regression (Fig. 7B, Supplementary Fig. S6A). H&E staining confirmed the profound reduction of metastatic tumour burden in Grn^−/−^ mice treated with αPD-1 compared to WT mice treated with αPD-1 (Fig. 7C). At endpoint, metastatic tumours in WT mice showed a marked accumulation of myofibroblasts, which was unaffected by αPD-1 therapy. As expected, lack of granulin resulted in a significant reduction of αSMA^+^ myofibroblast accumulation independent of αPD-1 treatment (Fig. 7D, E). In WT tumours, we found few numbers of CD8^+^ T cells within metastatic lesions of control IgG treated tumours, and CD8^+^ T cells numbers remained unaffected by αPD-1 administration (Fig. 7F). In contrast, depletion of granulin caused a significant increase in CD8^+^ T cell numbers in tumours in response to αPD-1 treatment (Figs. 7D, F). Next we assessed the activation state of CD8^+^ T cells by measuring GzmB and IFNγ expression levels in CD8^+^ T cells isolated from metastatic tumours. While αPD-1 treatment did not alter GzmB nor IFNγ expression levels in CD8^+^ T cells isolated from WT tumours, depletion of granulin led to a significant upregulation of GzmB and IFNγ expression in CD8^+^ T cells in response to αPD-1 administration (Figs. 7G, H).

**Figure 7:**
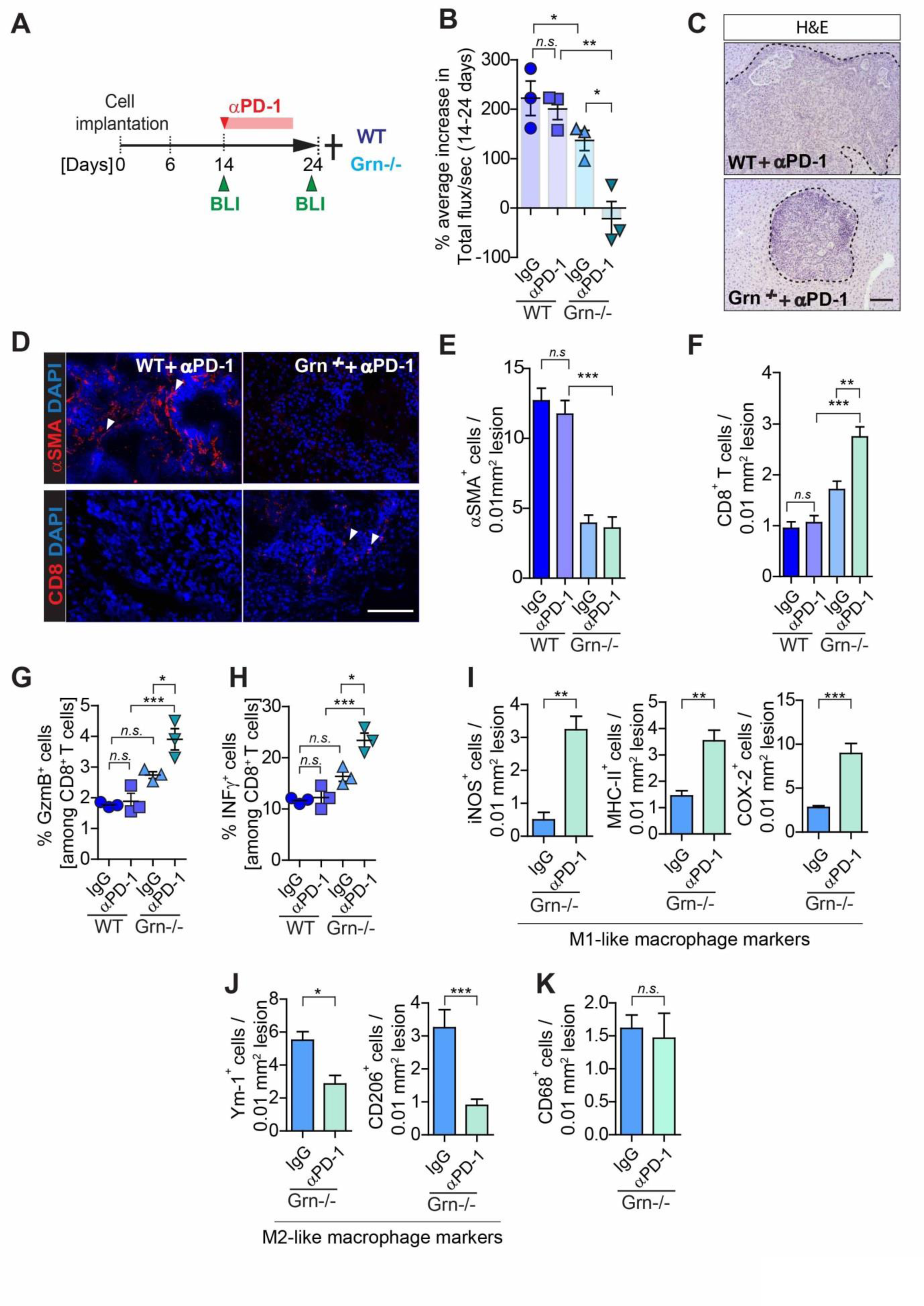
*Depletion of granulin restores response of metastatic PDAC to PD-1 therapy*. (**A-K**) Liver metastasis was induced by intrasplenic implantation of KPC^zsGreen/luc^ cells in WT and Grn^-/-^ mice and initial tumour burden was quantified on day 14 by *in vivo* bioluminescence imaging (BLI). On the same day, treatment regimen for control IgG or anti-PD-1 Abs was started. At day 24, change in tumour burden was quantified by *in vivo* BLI imaging and livers were extracted for analysis (n = 3 mice/group). (**A**) Schematic illustration of the experiment. (**B**) Percentage of average change in metastatic tumour burden (total flux/sec) in response to treatment assessed by *BLI* (individual data points, horizontal lines represent mean ± SEM). (**C**) Representative images of H&E staining of anti-PD-1 treated cohorts. Metastatic lesions are delineated with dashed lines. (**D**) Representative immunofluorescence images of myofibroblasts (αSMA^+^) and CD8^+^ T cell stainings of sequential liver sections from WT and Grn^−/−^ mice treated with anti-PD-1. (**E-F**) Quantification of myofibroblasts (αSMA^+^) accumulation and CD8^+^ T cell infiltration as described in D. (n = 3 mice / group; mean ± SEM) (**G-H**) Flow cytometry analysis quantifications of (G) Granzyme B (GzmB) and (H) IFNγ^+^ cells among metastasis infiltrating CD8^+^ T cells (n = 3 mice /group, individual data points, horizontal lines represent mean ± SEM). (**I-K**) Quantification of the number of cells staining positive for (I) pro-inflammatory M1 -like (iNOS, MHC-II, COX-2) MAMs, (J) anti-inflammatory M2-like (Ym-1, CD206) MAMs, and (K) total (CD68) MAMs markers in hepatic metastatic lesions from (A) based on IHC analysis (n=3 mice/group). Scale bar = 100 μm; *P*<0.001; **, *P* < 0.01; *, *P* < 0.05; n.s, not significant.

We next queried whether the observed increase in IFNγ expression by CD8^+^ T cells in αPD-1 treated granulin deficient mice promotes an immune stimulatory M1 -like MAM phenotype, allowing them to further fuel an anti-tumour immune attack. Indeed, we found in granulin deficient tumours treated with αPD-1 an increased presence of immune stimulatory MAMs that expressed pro-inflammatory markers, iNOS, MHC-II, and COX-2 (Fig. 7I; Supplementary Fig. S6C), while total macrophage numbers remain the same (Fig. 7K; Supplementary Fig. S6B). Reciprocally, these tumours displayed lower numbers of immunosuppressive Ym-1^+^ and CD206^+^ MAMs compared to untreated Grn^−/−^ tumours (Fig. 7J; Supplementary Fig. S6D).

Although both CSF-1 inhibition and granulin depletion are able to restore cytotoxic T cell infiltration in metastatic tumours, we observed that αPD-1 treatment was more effective in granulin deficient tumours (Fig. 7B) compared to tumours treated with αCSF-1 (Fig. 4B). Thus, these tumours might still be immunologically different and respond differently to immunotherapy. A recent study revealed that blockade of CSF-1R expressed on cancer associated fibroblasts can induce the accumulation of immunosuppressive Ly6G^^+^^ cells, thereby counteracting the therapeutic benefit of CSF-1R and/or αPD-1 inhibition (44). In agreement with these studies, we found that αCSF-1 treatment increases the infiltration of Ly6G^+^ cells in metastatic tumours, while granulin depletion did not have this effect (Supplementary Fig. S6E).

Taken together, these findings demonstrate that depletion of granulin dramatically improves the response of metastatic pancreatic tumors to αPD-1 treatment *in vivo* and that targeting a macrophage secreted factor such as granulin might be more effective than targeting macrophages.

## Discussion

The data presented herein describe that macrophage-derived granulin is a critical inducer of T cell exclusion in metastatic PDAC, and provide pre-clinical data that support the rational for targeting granulin in combination with immune checkpoint blocker PD-1 for the treatment of metastatic PDAC (Supplementary Fig. S7).

During cancer progression, immune evasion is a fundamental mechanism which allows malignant tumour cells to escape destruction by the effector cells of our immune system (1). Pancreatic cancer is a highly aggressive metastatic disease, yet little is known about the composition of the TME at the metastatic site and its role in protecting disseminated cancer cells from an immune response (18). A recent study described how the presence of tumour infiltrating CD8^+^ T cells together with the presence of neoantigen numbers stratifies PDAC patients with the longest survival; emphasizing the critical role of CD8^+^ T cell in inhibiting PDAC progression (45). In agreement with these findings, we observed that CD8^+^ T cell function and infiltration are lost during metastatic progression. We found that small hepatic metastatic PDAC deposits were well infiltrated by cytotoxic CD8^+^ T cells, which appeared partially functional, based on the elevated expression levels of activation markers and the observed increase in apoptosis of disseminated cancer cells. In contrast, during the progression into larger metastatic deposits, T cell infiltration is lost, and cancer cells survive. The loss of cytotoxic T cell function and infiltration correlates with an increase of M2-like MAMs in PDAC metastasis. Macrophages demonstrate a high degree of plasticity, and can be polarized into M1 -like immune stimulatory and M2-like immunosuppressive macrophages (40). We found an abundant number of M1-like MAMs surrounding small metastatic deposits. Transcriptional gene expression analysis of tumour derived MAMs confirmed their immunostimulatory phenotype. In contrast, large metastatic deposits were surrounded by M2-like MAMs, which revealed an upregulation of immunosuppressive factors and showed potent CD8^+^ T cell suppressive functions, indicating that an immunosuppressive TME forms during hepatic metastatic PDAC progression.

The importance of targeting the immunosuppressive TME in PDAC to obtain clinical benefit from immunotherapy is becoming increasingly more evident (46). Unfortunately, immune checkpoint blockade monotherapies in PDAC have shown limited success (16,17) and it was proposed to be caused by the presence of a rich immunosuppressive TME found in the affected pancreas that unables T cell infiltration and function (47). Based on our findings, we explored the therapeutic use of the immune checkpoint inhibitor αPD-1 alone or in combination with therapies targeting MAMs or their function. Single agent αPD-1 therapy instigated during early stages of metastatic development was efficient in enhancing cytotoxic CD8^+^ T cell activity, and in reducing metastatic progression. However, αPD-1 therapeutic benefits were completely ablated when PD-1 antagonist were administered in mice harbouring advanced metastatic lesions. Noteworthy, central to the efficacy of immune checkpoint blockade is the requirement for cytotoxic CD8^+^ T cells to infiltrate into tumours (48). Indeed, we found that metastatic tumours responding to αPD-1 treatment revealed abundant T cell infiltration within tumour lesions, while, in αPD-1 resistant tumours, T cells accumulated in αSMA^+^ rich regions surrounding the metastatic lesions. Interestingly, the administration of αCSF-1 reduced the overall numbers of MAMs and favoured an immunostimulatory M1-like phenotype, and also reduced the desmoplastic reaction at the metastatic site, which was accompanied by a marked increase in cytotoxic CD8^+^ T cell infiltration into tumours. In PDAC, dense fibrosis and myofibroblast activation have been recognized to be a barrier for chemotherapy and T cell infiltration and are now recognized as regulators of immune surveillance and immunotherapy efficacy for primary pancreatic tumours (20,47). However, the composition of the TME and its role in the immune response in metastatic pancreatic cancer has not yet been explored. In our studies we find that reduced fibrosis increases cytotoxic T cell infiltration and sensitizes metastatic lesions to αPD-1 treatment.

In respect to the anti-tumorigenic effect of targeting MAMs by blocking the CSF-1/CSF-1R signalling axis as monotherapy, we found that pharmacological blockade of CSF-1 by neutralizing antibody or inhibition of CSF-1R by small molecule inhibitor both reduced MAM numbers, and abolished the reprogramming of remaining MAMs towards an immunosuppressive M2-like phenotype when administrated during an early stage of PDAC metastasis. Accordingly, early interference with CSF-1/CSF-1R signalling led to an improved anti-tumour immune reaction at the metastatic site since CD8^+^ T cells isolated from metastatic lesions displayed significantly higher activation, together with a reduction of metastatic burden in a CD8^+^ T cell dependent manner. Noteworthy, a recent study revealed an increase in immunosuppressive Ly6G^+^ cells accumulation at the tumour site in response to blocking CSF-1R expressed on cancer associated fibroblasts, thereby counteracting its therapeutic benefits (44). In agreement with this study, we also found that αCSF-1 treatment increases the infiltration of Ly6G^+^ cells in established PDAC metastatic lesions. These observations suggest that hepatic metastatic tumours might counteract αCSF-1R macrophage based therapies by compensatory infiltration of other immune suppressive cell populations (44).

Dependent on the microenvironmental cytokine milieu, macrophages are not only promoting tumour growth, but they can also critically orchestrate an anti-tumour immune response (40). Thus, therapies that aim to specifically inhibit the pro-tumorigenic functions of macrophages, while sparing and/or enhancing their tumoricidal activity, could act as an alternative, and perhaps, more efficient approach than therapies that reduce macrophage numbers in tumours (49,50). In this regard, our studies indicate that depletion of granulin, a macrophage-derived factor that promotes transactivation of hepatic stellate cells into myofibroblasts (11), is sufficient to restore T cell infiltration at the metastatic site. Importantly, granulin deficient mice revealed a superior response to αPD-1 treatment compared to wildtype mice, with a marked increase of cytotoxic CD8^+^ T cell activation. The anti-tumorigenic effect of αPD-1 administration to granulin deficient mice was also superior compared to αPD-1 administration in combination with αCSF-1 macrophage-targeted therapy. αCSF-1 treatment also reduced MAM numbers at the metastatic site and induced a compensatory infiltration of potentially immunosuppressive Ly6G^+^ cells, while granulin depletion did not. Thus, in the presence of αPD-1, the immune system in granulin deficient mice can access an abundant numbers of macrophages for their reprogramming towards an immune stimulatory M1 -like phenotype, which facilitates the mounting of an effective immune response against cancer.

In conclusion, our studies show that in metastatic PDAC, granulin secretion by macrophages is induced by CSF-1 and prevents cytotoxic CD8^+^ T cell infiltration into metastatic lesions. Depleting granulin restores T cell infiltration and renders metastatic PDAC responsive to αPD1 therapy (Supplementary Fig. S7). These studies uncover a mechanism by which metastatic PDAC tumours evade the immune response and provide the rationale for targeting granulin and other factors promoting hepatic fibrosis, in combination with immune checkpoint inhibitors for the treatment of metastatic PDAC.

## Acknowledgements

We thank A. Santos, and L. Ireland for technical assistance. We thank the flow cytometry and cell sorting facility and the animal facility at the University of Liverpool for provision of equipment and technical assistance. We are grateful to H. Poptani and A. Minhas, Centre for Pre-Clinical Imaging, University of Liverpool, for their assistance with MR imaging. We acknowledge the Liverpool Tissue Bank for provision of tissue samples. We thank L. Young, CR-UK Cambridge Research Institute, for assistance with animal models. We also thank the patients and their families, who contributed tissue sample donations to these studies. These studies were supported by grants from the Medical Research Council (grant numbers MR/L000512/1 and MR/ /P018920/1) and the Pancreatic Cancer Research Fund (M.C.S.), North West Cancer Research Doctoral Training Programme (V.Q.), a National Institute for Health Research Biomedical Research Unit funding scheme through a NIHR Pancreas BRU/Cancer Research UK PhD fellowship (S.R.N.), and a Sir Henry Dale Fellowship jointly funded by the Wellcome Trust and the Royal Society (A.M., grant number 102521/Z/13/Z).

## Author contributions

V.Q. performed most of the experiments, designed experiments, analysed the data, and contributed to the preparation of the manuscript. C.R. and S.R.N helped with *in vivo* experiments. M.R. helped with tissue section analysis, *in vitro* assays, and gene and protein expression analysis. V.Q. and M.S.A. performed functional T cell assays. C.R. and V.Q performed flow cytometry and cell sorting. D.E. and D.T. provided primary murine KPC-derived pancreatic cancer cells. A. T. and T.M. provided lentiviral constructs and performed transduction of KPC-derived and Panc02 cells. F.C. and D.P. provided patient samples and helped with the analysis and interpretation of tumour biopsies. A.M. provided conceptual advice, designed experiments, and wrote the manuscript. M.C.S. conceived and supervised the project, interpreted data, and wrote the manuscript. All authors critically analysed and approved the manuscript.

